# Retinoic acid rewires the adrenergic core regulatory circuitry of childhood neuroblastoma

**DOI:** 10.1101/2020.07.23.218834

**Authors:** Mark W. Zimmerman, Adam D. Durbin, Shuning He, Felix Oppel, Hui Shi, Ting Tao, Zhaodong Li, Alla Berezovskaya, Yu Liu, Jinghui Zhang, Richard A. Young, Brian J. Abraham, A. Thomas Look

**Affiliations:** Department of Pediatric Oncology, Dana-Farber Cancer Institute, Boston, MA, 02215; Division of Molecular Oncology, Department of Oncology, St. Jude Children’s Research Hospital, Memphis, TN 38105; College of Animal Sciences, Zhejiang University, Hangzhou, Zhejiang 310052, China; National Clinical Research Center for Child Health, National Children’s Regional Medical Center, Children’s Hospital, Zhejiang University School of Medicine, Hangzhou, Zhejiang 310052, China; Cancer Center, Zhejiang University, Hangzhou, Zhejiang 310058, China; Department of Computational Biology, St. Jude Children’s Research Hospital, Memphis, TN 38105; Whitehead Institute, Cambridge, MA, 02142; Department of Biology, Massachusetts Institute of Technology, Cambridge, MA, 02142

**Keywords:** neuroblastoma, transcription, cell state, enhancer hijacking

## Abstract

Neuroblastoma cell identity depends on a core regulatory circuit (CRC) of transcription factors that collaborate with *MYCN* to drive the oncogenic gene expression program. For neuroblastomas dependent on the adrenergic CRC, treatment with retinoids can inhibit cell growth and induce differentiation. Here we show that when *MYCN*-amplified neuroblastomas cells are treated with retinoic acid, histone H3K27 acetylation and methylation become redistributed to decommission super-enhancers driving the expression of *PHOX2B* and *GATA3*, together with the activation of new super-enhancers that drive high levels of *MEIS1* and *SOX4* expression. These findings indicate that treatment with retinoids can reprogram the enhancer landscape, resulting in downregulation of *MYCN* expression, while establishing a new retino-sympathetic CRC that causes proliferative arrest and sympathetic differentiation. Thus, we provide mechanisms that account for the beneficial effects of retinoids in high-risk neuroblastoma and explain the rapid downregulation of expression of *MYCN* despite massive levels of amplification of this gene.

## Introduction

Cell identity is established by transcriptional core regulatory circuits (CRCs) composed of specific transcription factors that are driven by super-enhancers and form interconnected autoregulatory loops that coordinately regulate gene expression to establish cell state (*1, 2*). Throughout development, cell multipotency and differentiation are controlled by gene expression programs that are hierarchically regulated by CRCs and their extended regulatory network (*3–5*). Lineage specification of the embryonic neural crest, in particular, is governed by a dynamic architecture of master transcription factors and regulatory networks that give rise to diverse cell lineages during development (*6*).

Pediatric neuroblastoma, a neural-crest-derived tumor of the peripheral sympathetic nervous system (*7*), arises most often in the adrenal medulla, where sympathetic progenitor cells can become transformed by aberrant expression of *MYCN* or *MYC* and fail to differentiate into mature sympathetic ganglia or neuroendocrine chromaffin cells (*8–11*). Neuroblastomas in patients and experimental cell lines generally possess one of two CRC modules – the immature neural crest-like or mesenchymal subtype, defined by high expression levels of the *PRRX1, YAP/TAZ* and *AP-1* transcription factor genes, or the more commonly observed adrenergic CRC, characterized by high expression levels of *HAND2, ISL1, PHOX2B, GATA3, TBX2*, and *ASCL1* (*12–14*). Current models suggest that the neural crest-derived progenitors normally give rise to committed progenitors with the adrenergic cell state, culminating in terminally differentiated sympathetic neuronal cells and chromaffin neuroendocrine cells of the peripheral sympathetic nervous system (*15, 16*). In neuroblastomas with *MYCN* amplification, this oncogene stabilizes the adrenergic CRC to drive the expression of its transcriptional regulatory network and enforce an immature neuroblast cell state, while suppressing developmental signals that would normally induce differentiation or senescence (*17*). However, *MYCN* overexpression also results in a vulnerability called transcriptional addiction and creates tumor-selective gene dependencies, which include the adrenergic CRC transcription factors that sustain high levels of *MYCN* gene expression (*18, 19*).

Retinoic acid-based therapeutics provide a clinical benefit for patients with neuroblastoma and other malignancies through their ability to suppress tumor growth and promote cell differentiation (*20–24*). Adrenergic neuroblastoma cell lines treated with retinoids frequently exhibit phenotypes associated with neuronal differentiation (*25, 26*). Hence, we hypothesized that pharmacologically induced differentiation of neuroblastoma cells with retinoids depends on reprogramming of the adrenergic CRC, leading to rapid downregulation of *MYCN* expression, even in the context of massive *MYCN* gene amplification. Using a combination of transcriptional and chromatin assays, we examined the regulation of adrenergic CRC transcription factors in response to retinoic acid treatment and show that this autoregulatory loop is reprogrammed into a “retino-sympathetic” CRC, resulting in rapid downregulation of *MYCN* expression coupled with the induction of cell differentiation and proliferative arrest.

## Results

### Retinoic acid treatment inhibits tumor growth in MYCN transgenic zebrafish

High-risk neuroblastoma patients receive 13-*cis* retinoic acid (13-*cis* RA, or isotretinoin) as a maintenance therapy following high-dose chemotherapy with autologous stem cell transplantation, but the mechanistic basis for the efficacy of retinoid treatment is not well understood (*7*). Using a faithful zebrafish model of *MYCN*-driven neuroblastoma (*dbh:MYCN*) (*9*), we tested the ability of isotretinoin to inhibit neuroblastoma tumor initiation and progression *in vivo*. Three-week-old transgenic zebrafish exhibiting green, GFP-fluorescent cell masses in the interrenal gland (analogous to the human adrenal medulla), were treated with DMSO or 2 μM 13-*cis* RA (Fig. 1a). Relative to the DMSO control group, zebrafish receiving isotretinoin had a median 70% reduction in adrenal size after 6 days of treatment (Fig. 1b,c).

**Figure 1.**
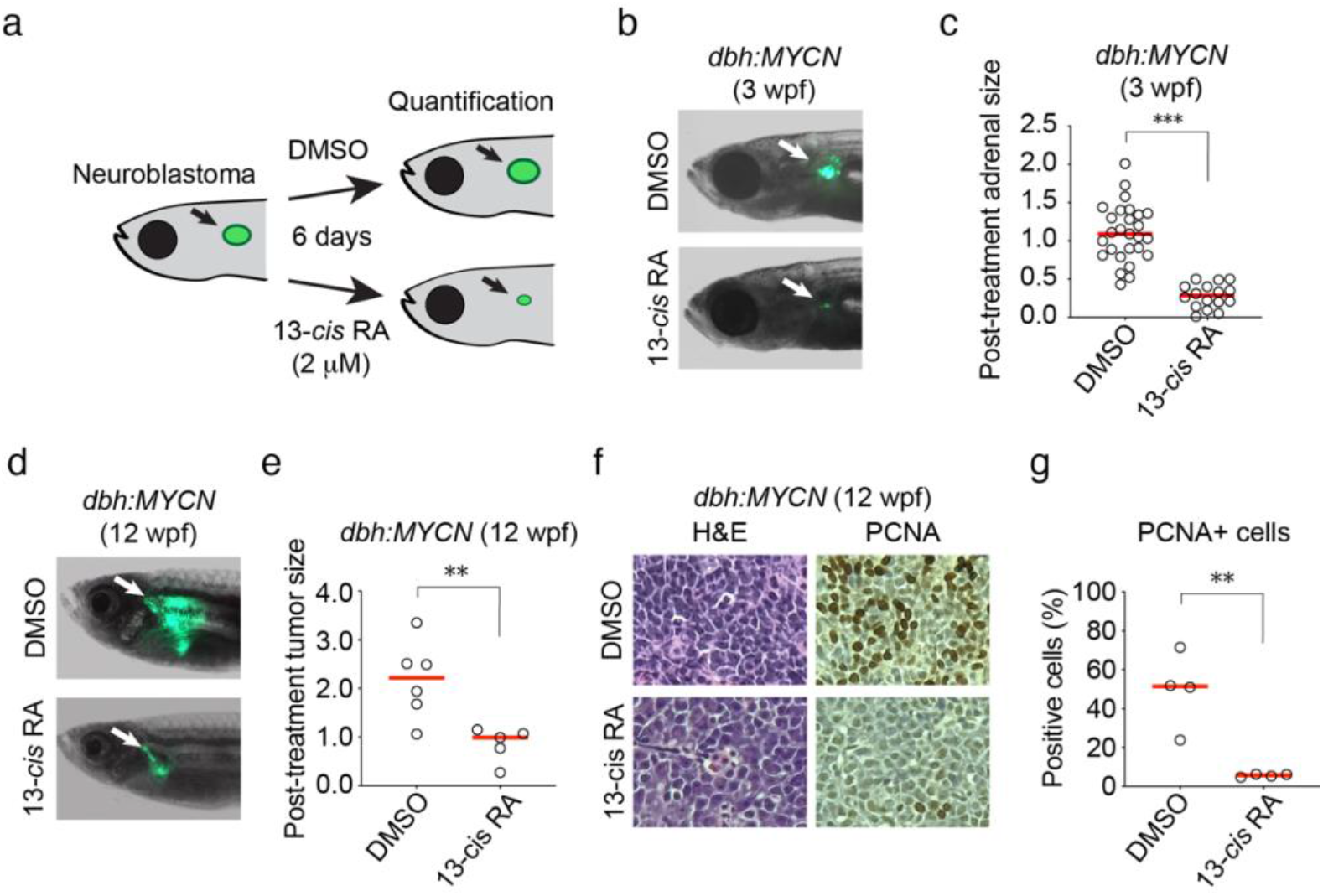
Retinoic acid suppresses the growth of transformed neuroblasts *in vivo*. a) Experimental design of the zebrafish tumor treatment assay. Tumor bearing *dbh:MYCN* transgenic zebrafish, 3 weeks post-fertilization (wpf), were treated with DMSO or 13-*cis* retinoic acid (2 μM) for 6 days. Tumor size was quantified by the EGFP+ surface area. b) Representative images of 3 wpf *dbh:MYCN* transgenic zebrafish with EGFP+ sympathoadrenal cell hyperplasia (white arrows) that were treated with DMSO or 2 μM 13-*cis* retinoic acid for 6 days. c) Dot plot showing the distribution of post-treatment tumor size of DMSO (n=26) or 13-*cis* retinoic acid (n=16) treatment in zebrafish, as quantified by EGFP cross sectional area. Red bar indicates the median value. ***p<0.001 by T-test. d) Representative images of 12 week-old *dbh:MYCN* transgenic zebrafish with EGFP+ neuroblastomas (white arrows) that were treated with DMSO or 5 μM 13-*cis* retinoic acid for 6 days. e) Dot plot showing the distribution of post-treatment tumor size of DMSO (n=6) or 13-*cis* retinoic acid (n=5) treatment in zebrafish, as quantified by EGFP cross sectional area. Red bar indicates mean. **p<0.01 by T-test. f) Representative images of 12 wpf neuroblastoma tumor sections treated with DMSO or 5 μM 13-*cis* retinoic acid for 6 days and stained with hematoxylin & eosin (left), or a PCNA detecting antibody counter stained with hematoxylin (right). g) Dot plot showing the percentage of PCNA+ cells in DMSO (n=4) or 13-*cis* retinoic acid (n=4) treated neuroblastoma tumors. Red bar indicates mean. **p<0.01 by T-test.

Next, *dbh:MYCN* transgenic zebrafish (12 wpf) were treated with DMSO or 5 μM 13-*cis* RA for 6 days. Similar to the treatment effect observed in juvenile zebrafish, mature zebrafish exhibited a 55% reduction in tumor burden (Fig. 1d,e). Histological analysis of DMSO and 13-*cis* RA treated tumor cells showed that the reduced tumor size was associated with loss of cell proliferation, as shown by a loss of PCNA staining (Fig. 1f,g, S1a,b). Additionally, the transcript levels of *raraa* and *rarab*, the two zebrafish orthologs of human *RARA*, were increased 3- to 5-fold in 13-*cis* RA treated tumors (Fig. S1c), which is similar to the effect of retinoic acid treatment on *RARA* expression in BE2C human neuroblastoma cells (Fig. S1d).

### ATRA treatment collapses the adrenergic CRC of MYCN-amplified neuroblastoma

To test the effects of retinoids on human *MYCN*-amplified neuroblastoma cells, we treated two *MYCN* gene amplified neuroblastoma cell lines (BE2C and NGP) with all-*trans* retinoic acid (ATRA), an active metabolite of isotretinoin, for 6 days and then examined its effects on cell growth and viability. BE2C and NGP cell growth was significantly suppressed by 6 days of 5 μM ATRA treatment relative to DMSO-treated control cells (Fig. 2a). Treatment with ATRA induced strong phenotypic changes in these cells, which included neurite outgrowth and upregulation of the structural proteins encoded by fibronectin (FN1), b3-tubulin (TUBB3) and vimentin (VIM) (Fig. 2b). These findings are consistent with previous studies reporting the ability of retinoids to induce differentiation in *MYCN*-transformed neuroblastoma cells (*27, 28*).

**Figure 2.**
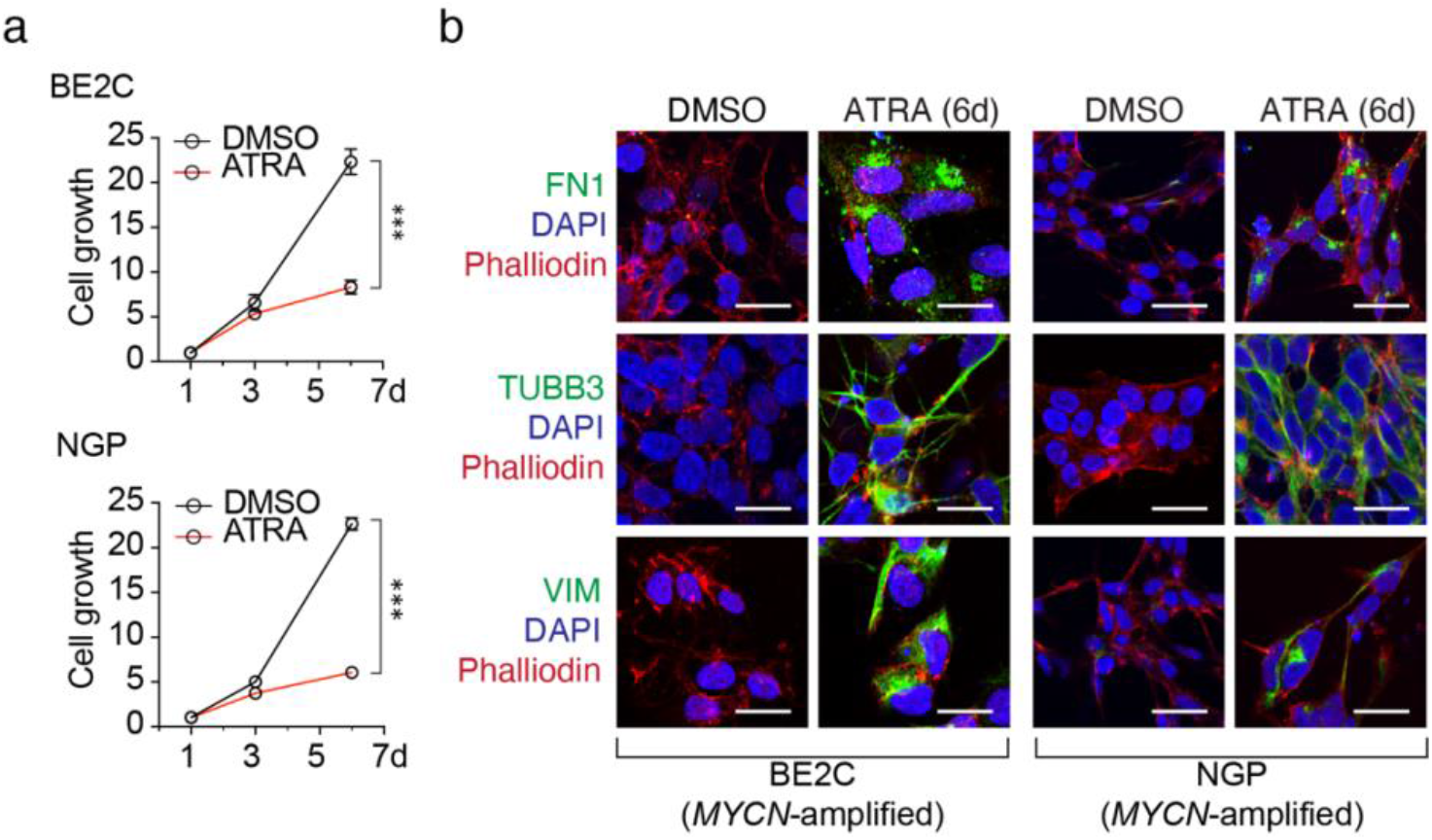
ATRA suppresses neuroblastoma cell growth and increases the expression of neuronal differentiation markers. a) Cell growth time course for BE2C and NGP cells comparing treatment with DMSO or 5 μM all-*trans* retinoic acid (ATRA) for 6 days; ***p<0.001 by ANOVA and T-test at 6 days. Data shows cell growth measurements with standard error bars for one representative experiment of three different independent experimental replicates. b) Confocal images of BE2C and NGP neuroblastoma cells treated with DMSO or 5 μM ATRA for 6 days. Cells were stained with fibronectin (FN1), b3-tubulin (TUBB3) or vimentin (VIM) (green) and counterstained with DAPI (blue) and phalloidin (red). Scale bar indicates 20 μm.

Cell state and fate specification depend on precisely controlled gene expression programs, which are often under the control of autoregulatory loops involving groups of key transcription factors, called the core regulatory circuitry or CRC (*2*). Thus, we investigated whether ATRA treatment affected the expression levels of members of the *MYCN*-driven adrenergic CRC and its extended regulatory network, which program the malignant cell state in most *MYCN*-amplified neuroblastoma cells (*19*). Using an ERCC spike-in normalized RNA-seq approach comparing cells after 6 days of treatment with either DMSO or ATRA, we observed dramatic changes in gene expression in ATRA-treated BE2C and NGP cells, with substantial downregulation of a subset of transcripts, including *MYCN, GATA3* and *PHOX2B* (Fig. 3a). Focusing on members of the adrenergic gene set described by Van Groningen *et al*. and Boeva *et al*. (*12, 13*), we noted downregulated expression of a subset adrenergic CRC transcription factors, including the members highly expressed in BE2C and NGP cells, several of which are tumor-selective gene dependencies (Fig. 3b) (*19*).

**Figure 3.**
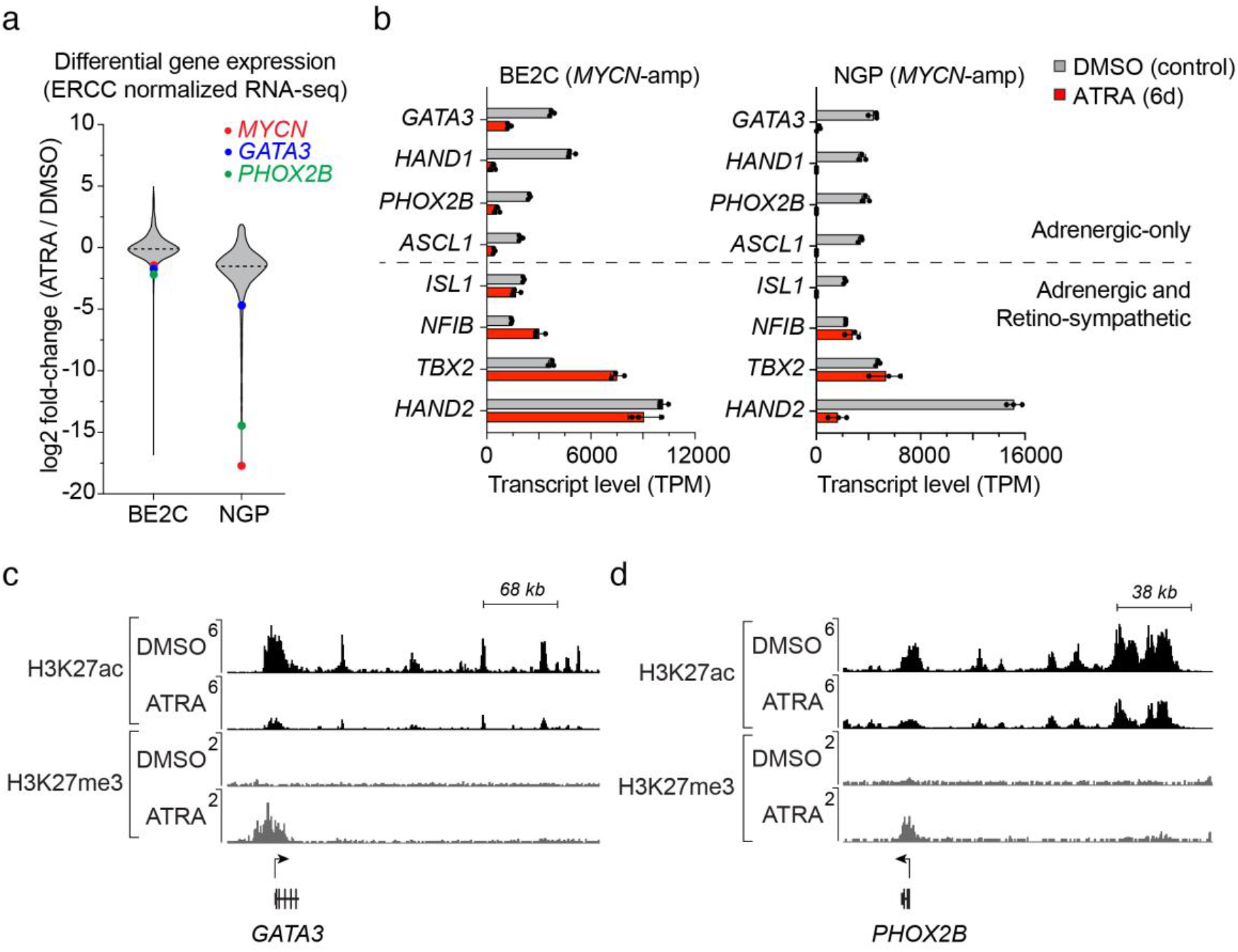
Effect of ATRA treatment on the expression and regulation of adrenergic CRC transcription factors in *MYCN*-amplified neuroblastoma. a) Violin plot illustrating log2-fold changes (ATRA/DMSO) among all highly expressed genes (base mean TPM > 10) when assayed by spike-in normalized RNA-seq in BE2C and NGP (*MYCN*-amplified) cells. Changes are highlighted for *MYCN* (red), *GATA3* (blue) and *PHOX2B* (green), all of which were reduced at the protein level as well. b) Expression levels of adrenergic CRC transcription factor genes determined by spike-in normalized mRNA-seq in BE2C and NGP cells treated with DMSO (grey) or 5 μM ATRA (red) for 6 days. c,d) Normalized ChIP-seq tracks for H3K27ac and H3K27me3 depicting super-enhancers associated with the *GATA3* (c) and *PHOX2B* (d) gene loci in BE2C cells. Cells were treated with 5 μM ATRA for 12 days. ChIP-seq read densities (y axis) were normalized to reads per million reads sequenced from each sample.

Some of the adrenergic transcription factors maintained high levels of expression despite a collapse of the adrenergic CRC (shown at the bottom of the bar graph in Fig 3b). This was unexpected because within a feed-forward autoregulatory loop, each transcription factor depends upon each of the others for its high levels of expression. We will demonstrate in the next section that this apparent paradox is explained because *TBX2, HAND2* and *ISL1* are retained as part of a new ATRA-driven CRC, called the retino-sympathetic CRC that forms as the adrenergic CRC collapses in ATRA-treated cells. Because CRC transcription factors bind coordinately within super-enhancers regulating down-stream genes within their extended regulatory networks, we examined genes regulated by ATRA-associated new super-enhancers that formed throughout the genome during ATRA treatment. We performed ChIP-seq for H3K27ac and H3K27me3 in *MYCN*-amplified neuroblastoma cells treated with DMSO or ATRA for 12 days and examined changes in the *cis*-regulatory regions associated with highly expressed genes. CRC transcription factors are associated with long stretches of H3K27ac-enriched chormatin, termed super-enhancers, that are capable of driving high levels of gene expression at their target promoters (*2, 29*). The *GATA3* and *PHOX2B* genes, which both showed reduced transcript levels after treatment with ATRA, lost H3K27ac enrichment in their associated enhancer regions and gained H3K27me3 modifications at their promoters, which are histone modification patterns associated with chromatin silencing (Fig. 3c,d) (*30*). Thus, ATRA-mediated differentiation of neuroblasts coincides with decreased expression of key CRC adrenergic transcription factors with corresponding changes in histone marks leading to repression of transcription.

### ATRA induces formation of a new CRC resulting in neuroblastoma cell differentiation

By contrast to the loss of H3K27ac associated with genes such as *GATA3* and *PHOX2B*, we also found that ATRA treatment induced enhancer signal increases, including the activation of super-enhancers associated with several retinoid-responsive genes (Fig. 4a,b, Fig. S2a,b). This enrichment occurred at discrete locations throughout the genome, and frequently resulted in the formation of new super-enhancers associated with several known retinoid-responsive genes, such as *CRABP2* and *EXOC6B*, whose expression was also increased following ATRA-treatment (Fig. S2c,d,e). Several transcription factor genes – including *SOX4* and *MEIS1* – acquired new super-enhancers as reflected by increased levels of H3K27ac modification in distal regulatory regions following ATRA treatment in both BE2C and NGP cells (Fig. 4c,d). This reflects the emergence of a new CRC induced by ATRA treatment that includes different transcription factors than untreated cells.

**Figure 4.**
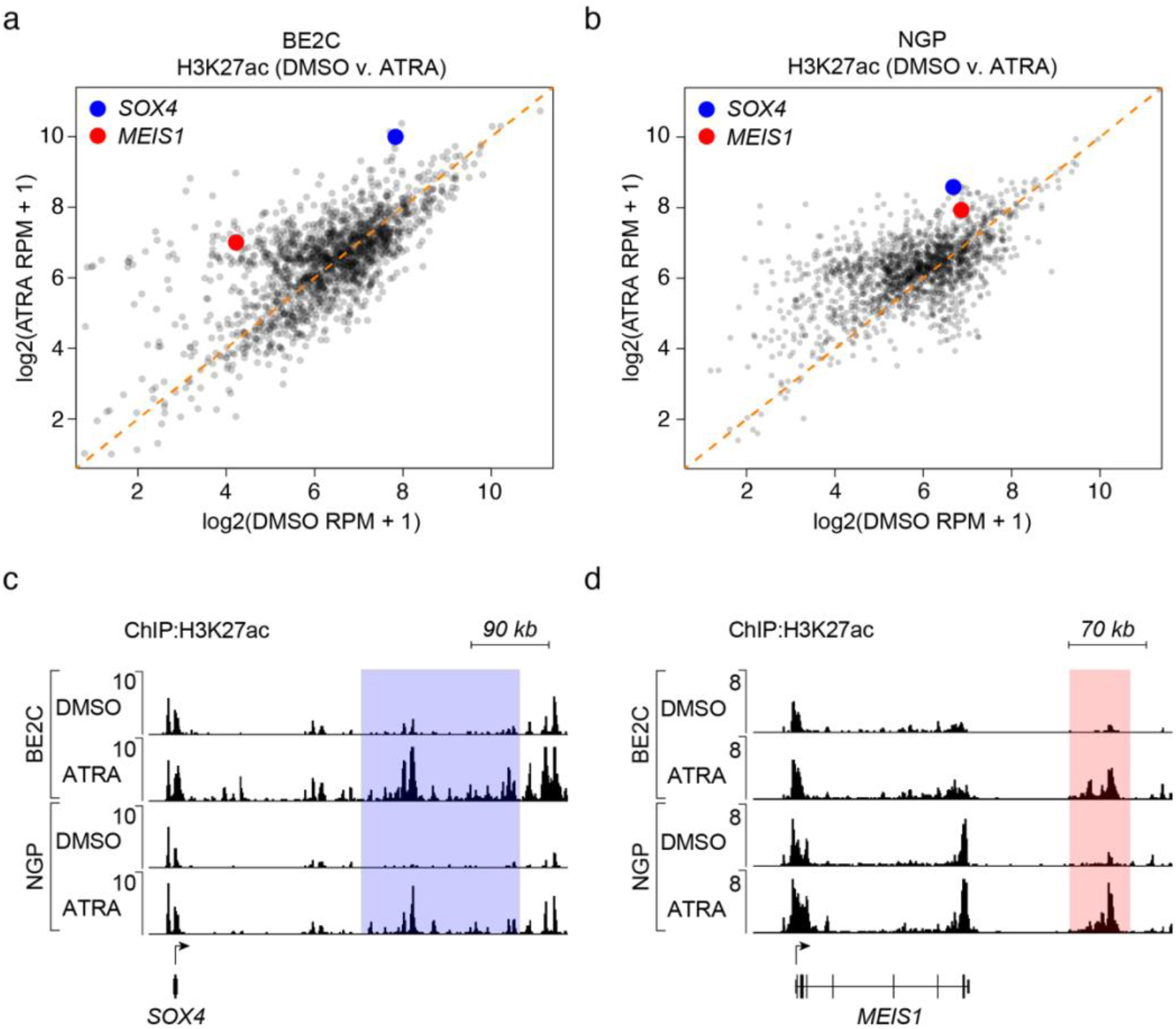
Treatment with ATRA redistributes H3K27ac modifications to remodel the enhancer landscape of neuroblastoma cells. a,b) Super-enhancers were identified in the DMSO and ATRA treated BE2C (a) and NGP (b) cells and collapsed into one set of regions whose differential enrichment was assessed in a H3K27ac coverage scatterplot. Orange diagonal line indicates equivalent H3K27ac signal in control DMSO compared to ATRA-treated cells. Highlighted enhancers were associated with SOX4 (blue) and MEIS1 (red). c,d) Normalized ChIP-seq tracks for H3K27ac showing acquired super-enhancers associated with SOX4 (c) and MEIS1 (d) in BE2C and NGP cells. Cells were treated with 5 μM ATRA for 12 days; shaded areas indicate super-enhancers identified by H3K27ac enrichment in the collapsed union list. ChIP-seq read densities (y axis) were normalized to reads per million reads sequenced from each sample.

To test the hypothesis that ATRA treatment induces a new CRC that replaces the adrenergic CRC, we examined the genes encoding transcription factors that are upregulated concomitantly with the activation of nearby super-enhancers that form after the start of ATRA treatment (Fig. 5a). We identified several genes encoding transcription factors that were highly expressed and associated with super-enhancers in BE2C and NGP cells that were treated with ATRA (Table S1). Because suitable antibodies are available for GATA3, PHOX2B, MEIS1 and SOX4, we performed CUT&RUN sequencing for enriched binding of these transcription factors in BE2C cells treated with DMSO or ATRA for 12 days. Comparing the enrichment of each of these transcription factors at the *HAND2* locus, which was highly acetylated under both treatment conditions, we note a shift from binding by GATA3 and PHOX2B in DMSO-treated cells, to predominant occupancy by MEIS1 and SOX4 within the same super-enhancer region in ATRA-treated cells (Fig. 5b,c). Additionally, comparing ChIP-seq results for binding by the RARA receptor, we show that RARA co-occupied enhancers with MEIS1 and SOX4 in ATRA-treated cells (Fig. 5c, Fig. S3). Similar binding patterns were observed genome-wide, where GATA3 and PHOX2B occupied super-enhancers in DMSO-treated cells, while MEIS1, SOX4 and RARA occupied super-enhancers in ATRA-treated cells (Fig. 5d).

**Figure 5.**
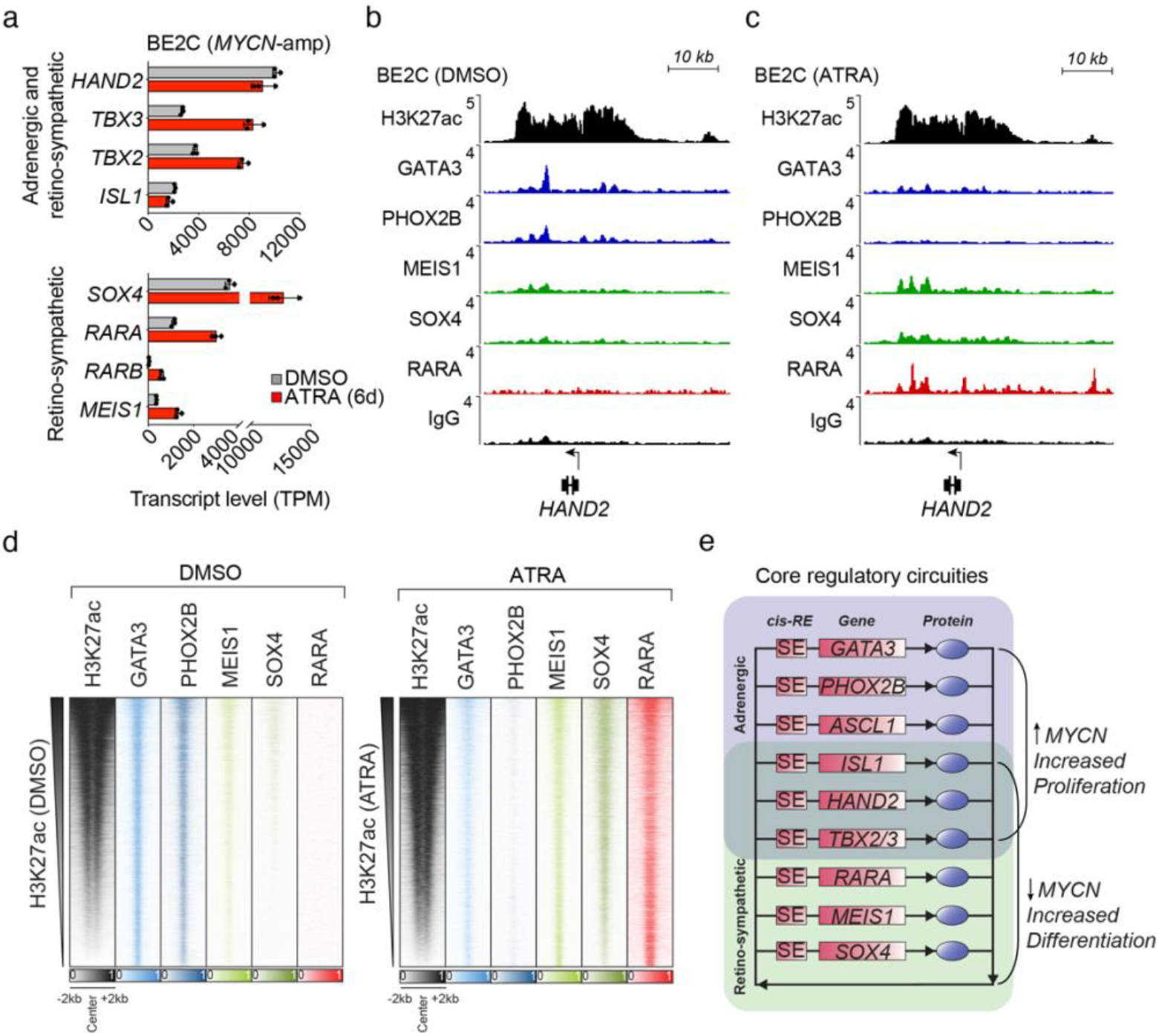
Cellular reprogramming by ATRA rewires neuroblastoma CRC favoring retino-sympathetic TFs. a) Transcript levels (TPM) of transcription factor genes upregulated in BE2C cells by treatment with ATRA (5 μM) for 6 days. A subset of transcription factors retained from the adrenergic CRC are shown above, and transcription factors unique to the retino-sympathetic CRC are shown below. b,c) Normalized ChIP-seq (H3K27ac and RARA) and CUT&RUN-seq (GATA3, PHOX2B, MEIS1 and SOX4) read coverage tracks depicting occupancy of transcription factors at the HAND2 gene locus in BE2C cells treated with DMSO (b) or ATRA (c) for 12 days. Rabbit IgG is shown as a control for the CUT&RUN-seq technique and alignment read densities (y axis) were normalized to reads per million reads sequenced. d) Genome-wide co-occupancy heatmap for adrenergic and retino-sympathetic transcription factors in DMSO- and ATRA-treated BE2C cells as determined by ChIP-seq (H3K27ac and RARA) and CUT&RUN-seq (GATA3, PHOX2B, MEIS1 and SOX4). Genomic regions (rows) were defined as those enriched in sequencing reads for at least one target and are ranked by the integrated H3K27ac signal. e) CRC transcription factors form an interconnected coregulatory loop, and treatment of adrenergic neuroblastoma cells with ATRA suppressed the expression and activity of *GATA3, PHOX2B* and *ASCL1*. Treatment with ATRA led to increased transcript levels and acquisition of new super-enhancers associated with *MEIS1* and *SOX4* in both BE2C and NGP cells. *RARA* had increased expression and were associated with super-enhancers under both DMSO and ATRA conditions. Regulatory elements and gene loci are denoted by rectangles, and proteins by oval symbols.

Treatment with ATRA has been reported to downregulate the expression of *MYCN* and induce cell cycle arrest with either differentiation or apoptosis in *MYCN*-amplified neuroblastoma cells (*27*). Our results indicate that the collective activity of the adrenergic CRC, including *HAND2, ISL1, PHOX2B, GATA3, ASCL1*, and *TBX2*, is essential for maintaining the high oncogenic expression level of *MYCN*, likely through activation of enhancers associated with the *MYCN* gene that are included in the amplified sequences (*31, 32*). Loss of *GATA3, PHOX2B* and *ASCL1* expression causes the adrenergic CRC to collapse after ATRA treatment, as noted in previous sections, accompanied by formation of the new retino-sympathetic CRC. Our results further indicate that the retino-*f*sympathetic CRC, which includes the transcription factors *RARA, SOX4* and *MEIS1*, as well as the shared members – *HAND2, ISL1* and *TBX2* – is not capable of activating native *MYCN* enhancers included within its amplicon, so *MYCN* levels are rapidly down-regulated despite the high levels of amplification of the gene (Fig. 5e, Fig. S4). *MYCN* is known to be a strong dependency factor in neuroblastomas with amplified *MYCN* (*19, 33*), accounting for the fact that neuroblastoma cells either undergo apoptosis or stop proliferating and undergo changes in gene expression consistent with terminal differentiation whenever the adrenergic CRC is dismantled and *MYCN* levels fall.

The transition from the adrenergic to the retino-sympathetic CRC in these cells required the continued presence of 5 μM ATRA to drive transcription through RAR/RXR. When ATRA was removed from the cells (washout), the retino-sympathetic CRC was lost, *MEIS1* and *SOX4* levels fell over 6 days, and reverted to baseline by 12 days after ATRA removal (Fig. S5). Concurrently the adrenergic CRC reformed and MYCN, PHOX2B and GATA3 levels returned to baseline over the 12 days after ATRA removal. The rapid reversal of cell state back to adrenergic after removing ATRA indicates that the neuronal differentiation during ATRA treatment is due to the regulatory activities of retino-sympathetic CRC transcription factors that induce changes in gene expression and epigenetic reprogramming, such as enhancer-activity modulation, rather than more permanent clonal alterations such as heritable DNA methylation.

### SOX4 and MEIS1 co-occupy their own and each other’s ATRA-driven super-enhancers

In addition to the ATRA-induced upregulation of expression and *de novo* formation of super-enhancers at *MEIS1* and *SOX4*, both of the encoded transcription factors co-occupy their own and each other’s super-enhancers, indicating an auto-regulatory expression loop (Fig. 6a,b). MEIS1 and SOX4 only lowly occupied their own and each other’s super-enhancers in the control condition, which reflects the low basal expression levels of each transcription factor prior to ATRA treatment. To assess mutual co-regulation between MEIS1 and SOX4 upon ATRA treatment, we disrupted the endogenous SOX4 gene using the CRISPR-Cas9 system. A time course experiment during 5 μM ATRA-treatment showed increased protein levels of SOX4 at 1, 2 and 3 days after treatment in control cells transduced with Cas9 and a non-targeting sgRNA (Fig. 6c). By contrast, the SOX4 protein was not detectable in BE2C cells transduced with Cas9 and a sgRNA targeting the *SOX4* coding region (*SOX4^-/-^*) at any point before or during ATRA treatment. Next, *MEIS1* gene expression was assayed at each time point in control and *SOX4^-/-^* cells by quantitative RT-PCR. Unlike control cells, in which *MEIS1* expression increased by 3.5-fold during ATRA treatment, *SOX4^-/-^* cells did not appreciably upregulate their expression level of *MEIS1* (Fig. 6d). Further, cells lacking SOX4 had lower expression levels of the differentiation-associated gene *FN1* during treatment with ATRA (Fig. 6e). Together, these results show that, as an integral member of the retino-sympathetic CRC, *SOX4* is essential for increased expression of the other retino-sympathetic CRC genes, as well as for downstream regulation of genes necessary for differentiation.

**Figure 6.**
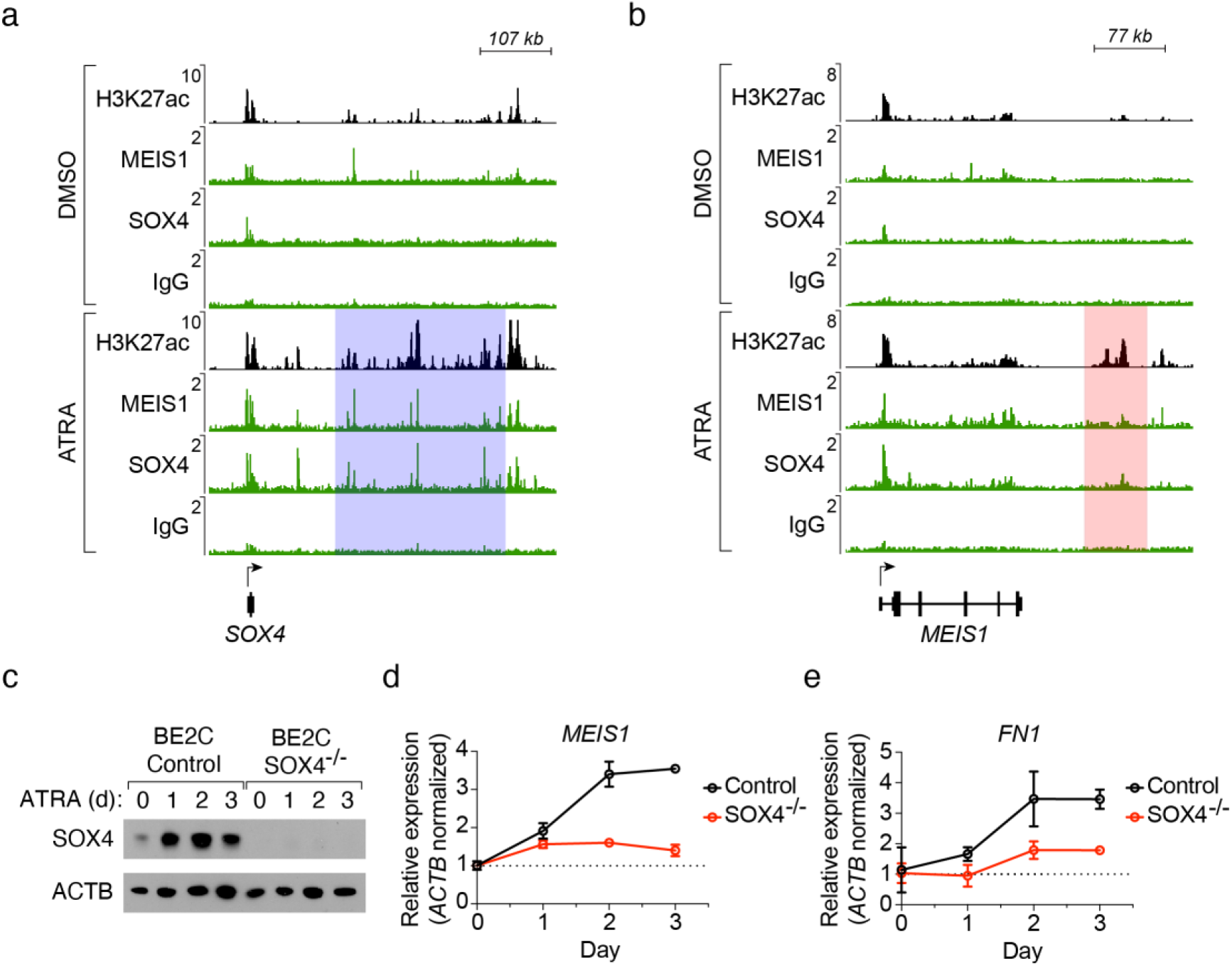
Co-regulated expression of *SOX4* and *MEIS1* mediates the shift from the adrenergic to retino-sympathetic gene expression program. a,b) Normalized ChIP-seq (H3K27ac) and CUT&RUN-seq (MEIS1, SOX4 and IgG) alignment tracks depicting occupancy of transcription factors at the SOX4 (a) and MEIS1 (b) gene loci in BE2C cells treated with DMSO (above) or ATRA (below) for 12 days. Super-enhancers were identified by H3K27ac signal and are shaded in ATRA-treated cells. Read densities (y axis) were normalized to reads per million sequenced from each sample. c) Western blot assay for SOX4 protein levels in wild-type control and *SOX4*-knockout BE2C cells treated with 5 μM ATRA for 0, 1, 2 and 3 days. ACTB was used as a loading control. d,e) Gene expression assayed by quantitative RT-PCR measuring the RNA levels of *MEIS1* (d) and *FN1* (e) in wildtype and *SOX4*-knockout BE2C cells treated with 5 mM ATRA for 0, 1, 2 and 3 days, and normalized to *ACTB*.

### Progressive waves of autoregulation establish the neuroblast differentiation program

Gene expression levels for the adrenergic neuroblastoma CRC members – *MYCN, HAND2, ISL1, PHOX2B, GATA3, ASCL1* and *TBX2* – were assayed by quantitative RT-PCR at 1, 3 and 6 days after treatment with 5 μM ATRA. *TBX2* steadily increases in expression levels, which is consistent with its joint membership in both the adrenergic and the new retino-sympathetic CRC (Fig. 7a,b). By contrast, the adrenergic CRC transcription factors that are not part of the retino-sympathetic *CRC – MYCN, GATA3, ASCL1* and *PHOX2B* – were quickly downregulated and expression levels stayed low during ATRA treatment (Fig. 7a). Two other adrenergic transcription factors, which are shared with the retino-sympathetic CRC – *HAND2* and *ISL1* – fell about 30-40 percent and then expression levels remained stable or slowly increased during 6 days of continuous treatment. By contrast, genes that do not belong to the adrenergic CRC and acquire new super-enhancers as they join the retino-sympathetic CRC induced by ATRA – *RARA, MEIS1, SOX4* – have steadily rising expression levels during ATRA treatment (Fig. 7b). Western blotting demonstrated that changes in expression levels for each of these transcription factors were concordant at the RNA and the protein levels obtained by Western blotting (Fig. 7c).

**Figure 7.**
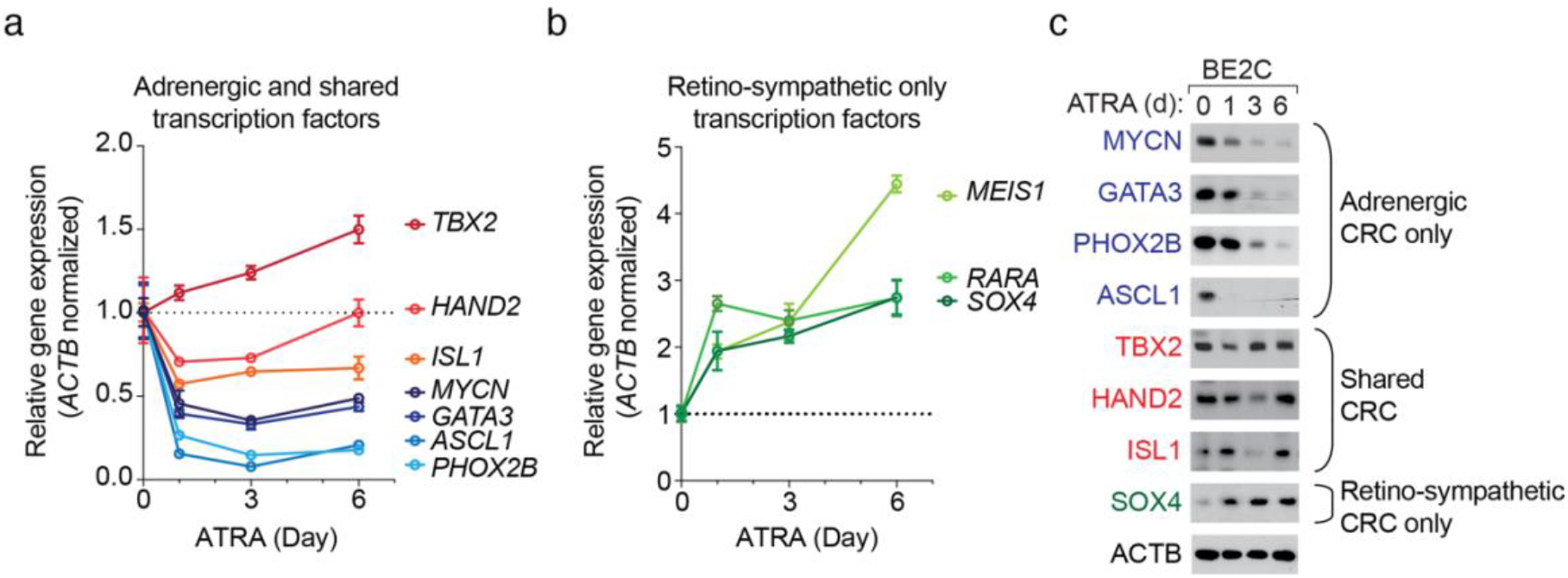
Rapid suppression of the adrenergic CRC and upregulation of the retino-sympathetic CRC when *MYCN*-amplified neuroblastoma cells are exposed to ATRA. BE2C cells were treated with 5 μM ATRA for 0, 1, 3 and 6 days, assayed by quantitative RT-PCR and normalized to *ACTB*. a) RNA gene expression levels of transcription factors exclusively in the adrenergic CRC, *MYCN, GATA3, PHOX2B*, and *ASCL1*, and those shared by both CRCs, *TBX2, HAND2, ISL1*. b) RNA gene expression levels of transcription factors exclusively in the retino-sympathetic transcription factors, *MEIS1, SOX4* and *RARA*. c) Western blot assay of protein levels for transcription factor belonging to either CRC in BE2C cells treated with 5 μM ATRA for 0, 1, 3 and 6 days. ACTB is shown as a protein loading control.

### ATRA resistance due to enhancer hijacking by the MYCN or MYC oncogenes

Previous studies have shown that downregulation of *MYCN* is a critical early event that is necessary to facilitate neuroblastoma cell differentiation in cells treated with ATRA (*33*). To demonstrate this effect experimentally, we transduced BE2C cells with an expression vector encoding *MYCN* or empty vector control. Western blotting demonstrated that this approach was able to enforce high levels of MYCN protein expression that was sustained in these cells despite suppression of endogenous MYCN during treatment with ATRA (Fig. S6a). To assess the induction of differentiation by ATRA, we examined expression of the retino-sympathetic target gene *FN1* by quantitative RT-PCR, a gene known to be expressed in ATRA-differentiated cells (*34*). We found that the expression of this gene was upregulated by ATRA in control cells, but that cells with sustained high levels of MYCN expression were unable to adopt the retino-sympathetic CRC and did not show increased *FN1* expression (Fig. S6b).

A subset of neuroblastoma cell lines that express high levels of *MYC* or *MYCN* do not respond to treatment with retinoids (*35–37*). One example is NBL-S, which expresses high levels of *MYCN* but does not have a high *MYCN* copy number. Instead, this cell line harbors a *t*(2:4) chromosomal translocation, which repositions the super-enhancer formerly associated with *HAND2* in close genomic proximity to *MYCN* on one allele – a phenomenon known as enhancer hijacking (Fig. 8a, Fig. S7). Similarly, the high *MYC*-expressing cell line SKNAS harbors a *t*(4:8) rearrangement, which repositions the *HAND2* super-enhancer to drive high levels of expression of the *MYC* gene (*11*). Expression of *HAND2* was not abolished in NBL-S by treatment with ATRA, and its associated super-enhancer remained stable when assayed by ChIP-seq (Fig. 8a). This outcome is consistent with the results in *MYCN*-amplified BE2C and NGP cells, where ATRA treatment had minimal effects on the stability of the super-enhancer regulating *HAND2* (see Fig. S4).

**Figure 8.**
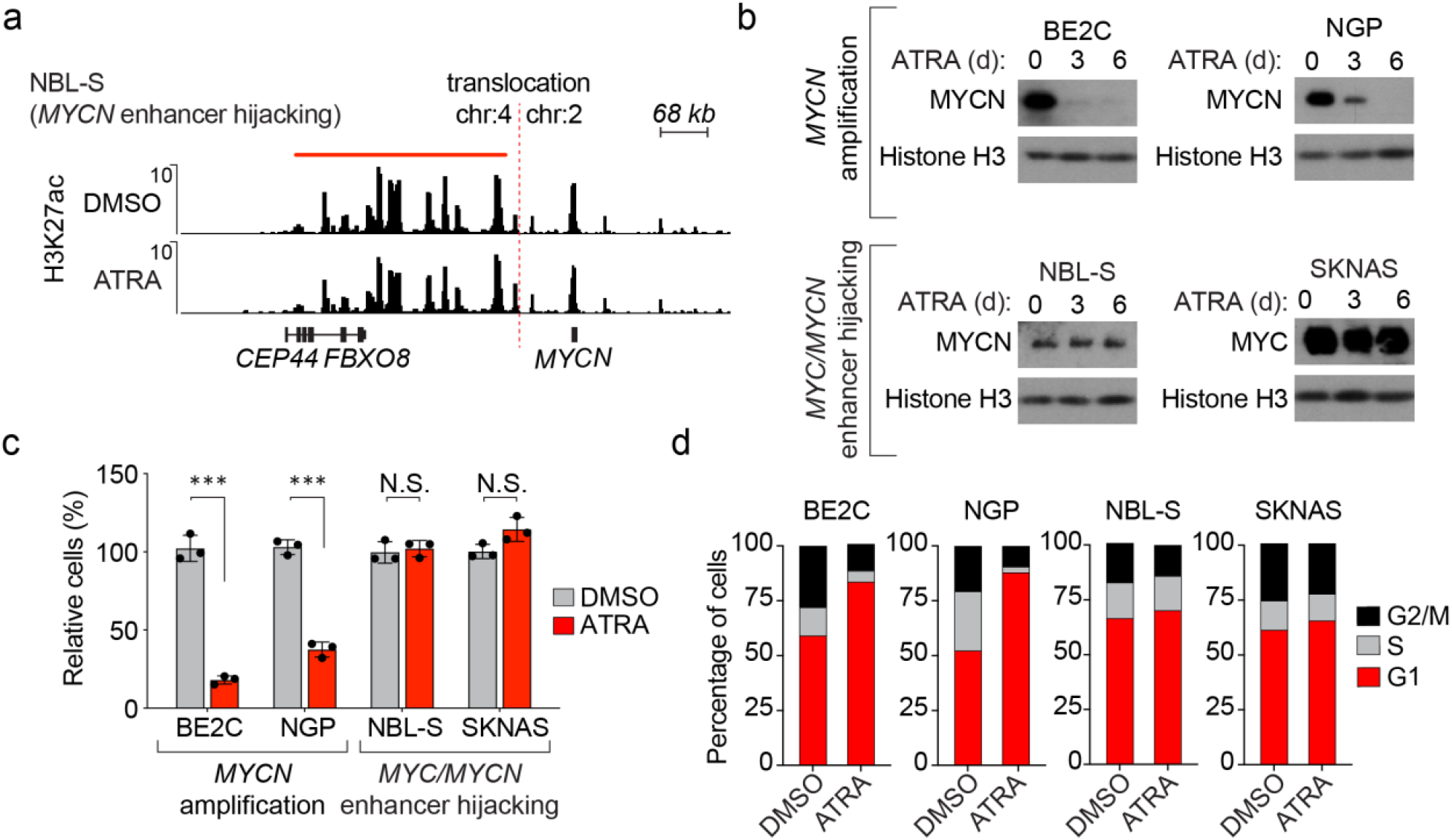
Hijacking the *HAND2* super-enhancer subverts ATRA-mediated suppression of *MYCN* or *MYC* expression. a) H3K27ac ChIP-seq in NBL-S cells treated with DMSO or ATRA (5 μM) for 12 days, showing the genomic region surrounding the *t*(2;4) structural variation involving the *MYCN* gene locus. The region downstream of *FBXO8* is a super-enhancer that is translocated to within close proximity of the *MYCN* gene. b) Western blot assay measuring MYCN or MYC protein levels in BE2C and NGP (*MYCN*-amplified), along with NBL-S and SKNAS (enhancer hijacking) cells following treatment with ATRA (5 μM) for 0, 3 and 6 days. c) Cell viability assay following 6 days of treatment with DMSO or ATRA (5 μM) in *MYCN*-amplified (BE2C and NGP) and *MYC/MYCN* enhancer hijacked (NBL-S and SKNAS) cell lines. ***p<0.001 by T-test; not significant (N.S.) d) Bar graphs showing cell cycle distribution determined from hypotonic citrate propidium iodide (PI) staining of BE2C, NGP, NBL-S and SKNAS cells treated with DMSO or ATRA (5 μM) for 6 days.

In BE2C and NGP cells, the MYCN protein level was almost completely depleted by 6 days of treatment with ATRA; however, NBL-S cells treated with ATRA showed sustained levels of MYCN protein at both 3 and 6 days (Fig. 8b). Thus, because the retino-sympathetic CRC binds and activates the *HAND2* super-enhancer, *MYCN* is still expressed at high levels in ATRA-treated NBL-S cells. These cells are blocked from differentiating and continue to proliferate as neuroblasts despite ATRA-induced activation of DNA binding by the RARA transcription factor. Several members of the retino-sympathetic CRC showed elevated transcript levels after ATRA treatment of these cells, but differentiation did not proceed because of sustained high levels of *MYCN* expression driven by the *HAND2* enhancer (Fig. S7). BE2C and NGP cells exhibited 72% and 63% reductions, respectively, in cell numbers when assayed 6 days after treatment with DMSO or ATRA (Fig. 8c). No significant difference in cell numbers was observed in DMSO and ATRA-treated NBL-S and SKNAS cells. Cell cycle phase distributions of propidium iodide-stained cells were analyzed by DNA flow cytometry before and after treatment with ATRA. By contrast to BE2C and NGP cells, which became blocked in G1 phase, NBL-S and SKNAS cells treated with ATRA continued to enter S phase and progress through the cell cycle, consistent with the continued cell proliferation by these *MYCN*- or *MYC*-hijacked neuroblastoma cell lines (Fig. 8d). Finally, we showed that depletion of the MYC protein in SKNAS with CRISPR-Cas9 was sufficient to sensitize these cells to treatment with ATRA and led to upregulation of retino-sympathetic target genes including *FN1* (Fig. S8). These results suggest that treatment strategies capable of attenuating MYC signaling could potentially be used in combination with ATRA to facilitate reprogramming.

## Discussion

13-*cis* retinoic acid is a clinically important component of current treatment protocols for pediatric neuroblastoma and has been shown to inhibit cell growth and induce differentiation when tested *in vitro* using many different neuroblastoma cell lines (*25, 28, 38–40*). We sought to explain why ATRA, the active metabolite of 13-*cis* retinoic acid, exerts these effects in neuroblastoma. Here we show that ATRA is capable of reprogramming the cell state of neuroblastoma, by fundamentally altering the core regulatory transcription factors that initiate and maintain the adrenergic gene expression program required for the tumorigenicity of these cells (*12, 13, 19*).

The adrenergic CRC consists of an interdependent autoregulatory network that includes *HAND2, ISL1, PHOX2B, GATA3, ASCL1* and *TBX2*, and is essential to drive the expression of *MYCN* and to facilitate the growth and survival of *MYCN*-amplified neuroblastoma cells (*14, 19*). Treating *MYCN*-amplified neuroblastoma cells with ATRA resulted in rapid downregulation of *MYCN* expression, loss of H3K27ac chromatin modifications associated with active super-enhancers within the *PHOX2B* and *GATA3* gene loci, and the acquisition of H3K27me3 chromatin silencing modifications of the promoters of these genes, which together lead to decreased expression levels of members of the adrenergic CRC. Despite the massively increased copy number of *MYCN* in these cells, *MYCN* expression levels were extremely sensitive to ATRA-treatment and became profoundly downregulated following collapse of the adrenergic CRC required to drive its expression. Thus, one effect of ATRA treatment is collapse of the adrenergic CRC due to direct or indirect repression of the *MYCN, GATA3, PHOX2B* and *ASCL1* genes. Although the transcript levels of these genes were dramatically reduced during the first day of ATRA-treatment, the MYCN, GATA3 and PHOX2B protein levels did not fall completely until up to 2 days later. This indicates that while the effects of ATRA on CRC gene RNA expression are immediate, transition to the new cell state requires additional time due to the delay in transcription factor protein turnover and new protein synthesis that is required for reprogramming of the transcriptome.

Our results demonstrate that the HAND2, ISL1 and TBX2 super-enhancers, along with high expression levels of their encoded mRNAs, were maintained after ATRA-mediated differentiation. Concomitantly, new super-enhancers were established at MEIS1 and SOX4, which coincided with increased expression levels of these genes. Thus, ATRA initiates a change in cell state of neuroblastoma cells corresponding to a shift from the adrenergic CRC to a new retino-sympathetic CRC, which includes the RARA, HAND2, ISL1, TBX2, TBX3, MEIS1 and SOX4 (Fig. 7 and Table S1). The genes of the retino-sympathetic CRC establish an extended regulatory network to enforce the differentiation of neuroblasts into mature sympathetic neuronal cells.

During ATRA-treatment, the mRNA levels for adrenergic CRC transcription factor genes that are not also components of the retino-sympathetic CRC – *MYCN, PHOX2B, GATA3*, and *ASCL1* – fell precipitously by day one, fell a little further by day 3, and then increased slightly by day 6 (Fig 7a). However, the corresponding protein levels decreased more slowly over the 6-day treatment period (Fig. 7c). Super-enhancers form nuclear condensates driven by the high concentration of transcription factors, co-factors and RNAs within a confined three-dimensional space in the nucleus (*29, 41*). It is possible that the changes in transcription during ATRA treatment are accompanied by a loss of CRC-mediated protection from ubiquitination such that these transcription factor proteins have reduced stability unless they are incorporated into a new CRC, as we observed for HAND2 and TBX2, which are retained in the retino-sympathetic CRC. Experiments to measure the protein half-lives of these classes of transcription factors during ATRA treatment would resolve this issue in the future.

Retinoids are vitamin A derivatives that have an essential function in vertebrate development by regulating gene expression programs, including a major role in specification of the nervous system (*42, 43*). Both isotretinoin and ATRA are capable of binding to RAR receptors (*44, 45*), including RARA, which is driven by a super-enhancer during ATRA treatment and is highly expressed as a functional member of the retino-sympathetic CRC. Upon treatment, ATRA binds to RARA, which binds to RXR, and the complexes bind to retinoid response elements coordinately with other members of the retino-sympathetic CRC to activate the expression of genes associated with neuroblastoma differentiation (*46*). Our findings indicate that ATRA-bound RARA acts as a potent activator and becomes an integral component of the retino-sympathetic CRC. Thus, the differentiation of neuroblastoma cells treated with ATRA likely reflects transcriptional reprogramming that occurs during normal PSNS development in response to endogenous retinoids during the maturation of migratory neuroblasts into non-proliferative sympathetic neurons and chromaffin cells (*43*).

Collapse of the adrenergic CRC mediated by ATRA includes marked downregulated expression of the amplified *MYCN* oncogene, whose expression must be reduced prior to neuroblast differentiation (*39*). This is achieved in neuroblastomas with *MYCN* gene amplification because endogenous *cis*-regulatory elements included in the *MYCN* amplicon are selectively activated by the adrenergic CRC (*31, 32*). This likely explains the physiologic expression of *MYCN* observed in non-transformed adrenergic neuroblasts that serve as the cell of origin for this from of neuroblastoma, and is supported by the theory that genomic amplification events require active gene expression in order to be selected for in tumorigenesis (*47*).

*MYCN* gene amplification is the most recurrent form of activation of a *MYC* family gene in neuroblastoma, but it is not the only mechanism. Neuroblastoma tumors can also upregulate the expression of *MYC* or *MYCN* by chromosomal structural rearrangements that hijack super-enhancers regulating the expression of *HAND2 (11)*, which is an important transcription factor in both the adrenergic and retino-sympathetic CRCs. Because *HAND2* is a member of both CRCs, the super-enhancer regulating it is active in both cell states. In neuroblasts with enhancer-hijacking, the super-enhancer on the intact allele drives *HAND2* expression while the HAND2 super-enhancer on the translocated allele continues to drive expression of *MYC* or *MYCN* as the retino-sympathetic CRC is attempting to form during ATRA treatment. Therefore, *MYCN* expression in these cells is not ablated by ATRA activating the retino-sympathetic CRC, preventing *MYCN* downregulation, which is essential for cell cycle arrest and differentiation (*28*). Thus, high levels of expression of either *MYC* or *MYCN* due to translocations hijacking next to the *HAND2* super-enhancer produce neuroblastoma cells that are resistant to the effects of ATRA in inducing neuroblastoma cell differentiation. Examples of this resistance phenotype include the cell lines NBL-S, with a t(2;4) activating expression of *MYCN*, and SKNAS, with a t(4;8) activating expression of *MYC*. In both cases, ATRA-treated cells retain high levels of *MYCN* or *MYC* oncogene expression at the RNA and protein levels and are blocked from undergoing differentiation. These findings highlight a resistance mechanism that could explain why some patients may not benefit and relapse, despite treatment of minimal residual disease with retinoids. Thus, our studies of transcriptional control of cell state in neuroblastoma not only provide insight into the role of ATRA in the treatment of high-risk neuroblastoma, but also reveal mechanisms that impart resistance to retinoids in some children with this disease.

## Materials and Methods

### Cell lines and proliferation assays

BE2C, SKNAS and 293T cells were purchased from ATCC; NGP and NBL-S cells were purchased from DSMZ. All neuroblastoma cell lines were cultured at 5% CO2 in RPMI media containing 10% FBS and 1% Penicillin-Streptomycin. Cells were routinely tested (every 3 months) for mycoplasma contamination and genotyped (every 12 months) by STR analysis at the Dana-Farber Molecular Diagnostic Core Facility. Cell proliferation was measured by plating 5000 cells per well in white 96-well plates in 100 uL of total media containing DMSO or 5 μM ATRA. Cell viability at each time point was assayed with Cell Titer glo (Promega) according to the manufacturer’s protocol.

### Lentiviral CRIPSR-Cas9 mutagenesis

Stable Cas9:sgRNA expressing cell lines were created using lentivirus produced in 293T cells. Briefly, sgRNA target sequences (Table S2) were cloned in the lentiCRISPRv2 vector (Addgene plasmid #52963) as previously reported (PMID:25075903). Plasmids were transfected using Fugene HD (Promega) along with pMD2.G (Addgene Plasmid#12259) and psPAX2 (Addgene plasmid #12260) to generate viral particles. Following lentiviral transduction, cells were selected with puromycin and expanded prior to evaluation.

### Compounds and reagents

Isotretinoin (13-*cis* retinoic acid) and ATRA (all-*trans* retinoic acid) were purchased from Selleckchem. Cell culture grade DMSO was purchased from ATCC. Compounds were resuspended in DMSO to a stock concentration of 100 mM and added directly to cell culture media or zebrafish water at the indicated concentrations.

### Zebrafish tumorigenesis assays

Transgenic zebrafish were developed as previously reported (*9*). All animal experiments were conducted at Dana-Farber Cancer Institute in accordance with animal care and use committee protocol #02-107. Wildtype and transgenic zebrafish were maintained under standard aquaculture conditions at the Dana-Farber Cancer Institute. Transgenic *dbh:MYCN* zebrafish were crossed to a stable *dbh:EGFP* expressing line and sorted for EGFP+ fluorescence. EGFP+ zebrafish 3 and 12 weeks-post fertilization (wpf) were treated with DMSO or 13-*cis* RA (2 μM for 3 wpf, and 5 μM for 12 wpf) added directly to the aquaculture water and refreshed daily. Zebrafish were imaged at day 0 prior to treatment and again following 6 days of exposure to either compound. Prior to imaging zebrafish were anesthetized with tricaine and subsequently monitored for neuroblastoma tumor progression. All comparative experimental groups for sympathoadrenal and neuroblastoma tissue quantification were imaged under the same conditions, and acquired fluorescent images were quantified using ImageJ software (NIH) by measuring the area of EGFP fluorescence. Overlays were created using ImageJ and Adobe Photoshop 7.0.1.

### Immunohistochemistry

Zebrafish for histological analysis were euthanized with tricaine, fixed in 4% paraformaldehyde at 4°C overnight and decalcified with 0.25 M EDTA (pH 8.0). Paraffin sectioning followed by hematoxylin and eosin (H&E) staining or IHC was performed at the Dana-Farber/Harvard Cancer Center Research Pathology Core. Primary antibody (PCNA, EMD Millipore, MAB424R, 1:100) binding was detected with the diaminobenzidine-peroxidase visualization system (EnVision+, Dako). Mayer’s hematoxylin was used for counterstaining. Slides were imaged using the Echo Revolve4 inverted microscopy system.

### Immunofluorescence and confocal microscopy

Cell were grown and compound treated on glass slides in 6-well cell culture plates. After 6 days of treatment with DMSO or ATRA, slides were incubated with a primary antibody at 4°C overnight (Table S3), washed with PBST, and then incubated with a secondary antibody for 2 hours at room temperature. Secondary antibodies were conjugated with Alexa Fluor 488 (Life Technologies). Alexa Fluor 568 Phalloidin (Life Technologies) and DAPI (BD Biosciences) were used for counter staining. Fluorescent images were taken with a Leica SP5X scanning confocal microscope at the Confocal and Light Microscopy core facility at Dana-Farber Cancer Institute.

### Western blotting

Protein samples were collected and lysed using RIPA buffer containing protease and phosphatase inhibitors (Cell Signaling Technology). Lysates were quantified by Bradford assay (Bio-rad), and 10 μg of extracted protein was separated using Novex SDS-PAGE reagents and transferred to nitrocellulose membranes (Life Technologies). Membranes were blocked in 5% milk protein and incubated with primary antibodies (Table S3) overnight followed by secondary HRP-linked goat anti-rabbit and anti-mouse (Cell Signaling) antibodies (1:1000) according to the manufacturers’ instructions. Antibody bound membranes were incubated with SuperSignal West Pico chemiluminescent substrate (Thermo-Fisher) and developed using HyBlot CL autoradiography film (Thomas Scientific). The antibodies used immunoblotting are listed in Table S3.

### Quantitative RT-PCR

Total RNA was harvested using the RNeasy kit (QIAgen) according to the manufacturer’s protocol. First-strand synthesis was performed with Superscript III (Invitrogen). Quantitative PCR analysis was conducted on the ViiA7 system (Life Technologies) with SYBR Green PCR Master Mix (Roche) using validated primers specific to each target each gene. Primer sequences are displayed in Supplementary Table S2.

### Spike-in normalized RNA-seq

DMSO and ATRA treated cells were grown in triplicate using 6-well plates and collected directly into Trizol. ERCC spike-in RNA was diluted 1:10 in nuclease-free water and added directly to Trizol lysates after being normalized to cell number as per the manufacturer’s protocol (Life Technologies). Libraries were prepared using Illumina TruSeq stranded specific sample preparation kits from 500ng of purified total RNA according to the manufacturer’s protocol. The finished dsDNA libraries were quantified by Qubit fluorometer (Thermo-Fisher), TapeStation 4200 (Agilent), and RT-qPCR using the Kapa Biosystems library quantification kit (Roche) according to manufacturer’s protocols. Indexed libraries were pooled in equimolar ratios and sequenced on an Illumina NextSeq 550 with single-end 75bp reads by the Dana-Farber Cancer Institute Molecular Biology Core Facilities. Reads were aligned to a reference genome containing the non-random chromosomes from hg19 and the sequences of ERCC probes using hisat2 with parameters --no-novel-juncs and -G set to a gene database file downloaded from RefSeq on 7/5/2017 to which positions of the ERCC probes were added. Coverage of the genes in this list was calculated using htseq-count with parameters -i gene_id --stranded=reverse -f bam -m intersection-strict. Violin plots were created using Prism 8.4.3 (GraphPad). Raw and processed data files were deposited to the NCBI GEO server under super-series GSE155002 (Table S4).

### CUT&RUN sequencing and initial processing

CUT&RUN coupled with high-throughput DNA sequencing was performed using antibodies listed in Table S3 and Cutana pA/G-MNase (Epicypher) according to the manufacturer’s protocol. Briefly, cells were washed and incubated with activated Concanavalin A beads for 10 min at room temperature. Cells were then resuspended in antibody buffer containing 0.01% digitonin, 1 mL of each antibody (Table S3) was added to individual cell aliquots and tubes were rotated at 4°C overnight. The following day, targeted chromatin digestion and release was performed with 2.5 mL Cutana pA/G-MNase and 100mM CaCl2. Retrieved genomic DNA was purified with the MinElute PCR purification kit and eluted in 10 mL of buffer EB. Sequencing libraries were prepared with the automated Swift 2S system, followed by 100bp-PE sequencing with Novaseq 6000.

Reads were aligned to the human reference genome (hg19) using bowtie v1.2.2 in single-end mode with parameters –k 2 –m 2 –best and –l set to the read length. For visualization, WIG files were created from aligned read positions using MACS v1.4 with parameters –w –S –space=50 –nomodel –shiftsize=200 to artificially extend reads to be 200bp and to calculate their density in 50bp bins. Read counts in 50bp bins were then normalized to the millions of mapped reads, giving reads per million (rpm) values. WIG files were visualized in the IGV browser version 2.7.2. Raw and processed data files were deposited to the NCBI GEO server under super-series GSE155002 (Table S4).

### ChIP-seq and initial processing

Chromatin Immunoprecipitation coupled with high-throughput DNA sequencing (ChIP-seq) was performed as previously described (*19*). The antibodies used for each experiment are listed in Table S3. For each ChIP, 5 μg of antibody coupled to 2 ug of magnetic Dynabeads (Life Technologies) were added to 3 mL of sonicated nuclear extract from formaldehyde fixed cells. Chromatin was immunoprecipitated overnight, crosslinks were reversed, and DNA was purified by precipitation with phenol:chloroform:isoamyl alcohol. DNA pellets were resuspended in 25 uL of TE buffer. Illumina sequencing, library construction and ChIP-seq analysis methods were previously described.

Reads were aligned to the human reference genome (hg19) using bowtie v1.2.2 with parameters –k 2 –m 2 –best and –l set to the read length. For visualization, WIG files were created from aligned read positions using MACS v1.4 with parameters –w –S –space=50 –nomodel –shiftsize=200 to artificially extend reads to be 200bp and to calculate their density in 50bp bins. Read counts in 50bp bins were then normalized to the millions of mapped reads, giving reads per million (rpm) values. WIG files were visualized in the IGV browser version 2.7.2. Raw and processed data files were deposited to the NCBI GEO server under super-series GSE155002 (Table S4).

### Super-enhancer Identification and Assignment

Super-enhancers in BE2C and NGP cells were separately identified using ROSE (https://bitbucket.org/young_computation/rose). Briefly, two sets of peaks of H3K27ac were identified using MACS with parameter sets –keep-dup=auto –p 1e-9 and –keep-dup=all –p 1e-9. Peaks identified that contact the region chr2:14817188-17228298 were discarded because they fall within the amplified genomic regions around MYCN. The collapsed union of regions called using both MACS parameter sets that do not contact the discarded MYCN-proximal region were used as input for ROSE with parameters -s 12500 -t 1000 -g hg19. Enhancers were assigned to the single expressed gene, which was defined as being in the top 2/3 of promoter (TSS +/- 500bp) H3K27ac coverage in a sample, whose transcription start site was nearest the center of the enhancer.

### Differential coverage analysis

ATRA-induced changes in H3K27ac coverage were assessed at the collapsed union of super-enhancers identified separately by ROSE in each of four samples (NGP/BE2C, DMSO/ATRA). Coverage in each region was assessed using bedtools intersect and normalized by dividing each value by the millions of mapped reads per sample. Collapsed super-enhancers were assigned to the nearest gene considered expressed in any of the four samples, where expression was defined as being in the top 2/3 of promoter (TSS +/-500bp) H3K27ac coverage.

### Cell cycle analysis

Cells were treated with DMSO or ATRA for 6 days in triplicate, and 500,000 cells per sample were collected and lysed in cold hypotonic propidium iodide (PI) / RNase solution (50 ug/mL PI, 4mM sodium citrate, 30 U/mL RNaseA, 0.1% TX-100). Samples were then vigorously vortexed and incubated at 37°C for 10 min. Sodium chloride was added to a final concentration of 0.15M, and stained nuclei were stored at 4°C until analysis was ready to be performed. The samples were then analyzed by flow cytometry (BD LSRFortessa). Cell cycle distribution was analyzed with the FlowJo cell cycle Watson (Pragmatic) model. The singlet population was isolated with a live cell gate. To analyze the proportion of cells in G1, S, and G2/M, the Watson (Pragmatic) model with the G2 peak constrained on G1 = G2 x 2 was used. Both debris and doublets were removed from the analysis.

### Statistical analysis

Statistical calculations were performed using Prism 7.01 (GraphPad). Digital images of the fluorescence signal for transgenic embryos, and the area of the fluorescence coverage, was quantified with ImageJ (NIH) for Fig. 1. Multivariate ANOVA analysis followed by two-tailed, unpaired t-tests with confidence intervals of 95% were used for the quantitative assays.

## Supplementary Materials

Figure S1. Loss of neuroblastoma cell proliferation following treatment with 13-cis retinoic acid.

Figure S2. Acquisition of new super-enhancers is associated with increased expression of their associated genes.

Figure S3. RARA and MEIS1 occupy H3K27ac-enriched super-enhancers associated with *MEIS1, HIC1* and *SOX4* in ATRA-treated neuroblastoma cells.

Figure S4. Several super-enhancers associated with CRC transcription factors are stable after treatment with ATRA.

Figure S5. Reversion to the adrenergic CRC and phenotype following ATRA washout.

Figure S6. Retained expression of MYCN blocks the induction of the ATRA-mediated differentiation program in neuroblastoma.

Figure S7. ATRA-mediated changes in gene expression and protein level in *MYCN*-amplified cells are not observed in cells that activate *MYCN* or *MYC* by enhancer hijacking.

Figure S8. Disruption of MYC activity sensitizes cells to the transcriptional effects of ATRA.

Table S1. Putative core regulatory transcription factors

Table S2. Oligo and primer sequences

Table S3. Antibody information

Table S4. NCBI GEO accession numbers

## Acknowledgments

We would like to thank J.R. Gilbert for editorial assistance and critical comments and Z. Herbert of the Dana-Farber Molecular Biology Core Facility for genomics support. This work was supported by NIH grants R35CA210064 (A.T.L.), R01CA216391 (J.Z.) and K08CA245251 (A.D.D), and the St. Jude Children’s Research Hospital Collaborative Research Consortium on Chromatin Regulation in Pediatric Cancer. M.W.Z. was supported by grants from the Alex’s Lemonade Stand Foundation, Charles A. King Trust, and Claudia Adams Barr Foundation. A.D.D. was supported by grants from the Alex’s Lemonade Stand Foundation, CureSearch for Children’s Cancer and Damon Runyon-Sohn Foundation. F.O. acknowledges the German Cancer Aid for their generous funding within the Mildred-Scheel-Postdoctoral program of the Mildred Scheel Foundation. A.D.D. and B.J.A. are supported by the American Lebanese Syrian Associated Charities (ALSAC).

## Author contributions

M.W.Z., A.D.D., S.H., R.A.Y., B.J.A. and A.T.L. conceived the project, performed data interpretation and wrote the manuscript with input from all authors. M.W.Z. and A.D.D. designed and performed ChIP-seq, RNA-seq and flow cytometry experiments. B.J.A. performed ChIP-seq and RNA-seq computational analysis. Y.L. and J.Z. analyzed neuroblastoma cell line genomic data. M.W.Z. and S.H. designed and performed in vivo zebrafish experiments. F.O., H.S. and T.T. performed cellular immunofluorescence assays and confocal microscopy. A.B. and Z.L. performed western blotting and other experiments.

## Data and materials availability

All data needed to evaluate the conclusions in the paper are present in the paper and/or the Supplementary Materials. ChIP-seq and RNA-seq data have been deposited in the NCBI Gene Expression Omnibus (accession number GSE155002).

